# Modeling Primary Fields of TMS Coils with the Fast Multipole Method

**DOI:** 10.1101/514919

**Authors:** Sergey N. Makarov, Lucia Navarro de Lara, Gregory M. Noetscher, Aapo Nummenmaa

## Abstract

In this study, an accurate TMS-coil modeling approach based on conductor’s cross-section representation with many distributed current filaments coupled with an efficient fast multipole method (FMM) accelerator is developed and tested. Uniform (Litz wire) or skin-effect based current distributions are included into consideration. Speed and accuracy estimates as well as two application examples are given, which indicate that this approach is potentially capable of rapid and accurate evaluation of various detailed TMS coil designs and arrays of such coils.

The MATLAB-based wire and CAD mesh generator for the coil geometry is interfaced with the FMM FORTAN program, which is also compiled within the MATLAB shell. No extra MATLAB toolboxes are necessary. The CAD model of the coil can be imported into any other computational software package in STL format. The algorithm is organized in the form of a MATLAB-based toolkit. First, a coil model is generated using a dedicated script. Then, we compute high-resolution 2D contour plots for any component of the electric and/or magnetic field in coronal, sagittal, and transverse planes via FMM. These two scripts may be further augmented with a parametric loop to enable rapid analysis.

**Objective:** We show how the fast multipole method can be applied to compute primary electric and magnetic fields for detailed TMS coil models

## 1. Introduction

For all three chief neurostimulation modalities – transcranial magnetic stimulation (TMS), transcranial electric stimulation (TES), and intracortical microstimulation (ICMS) – modeling the electric fields within a patient-specific head model is the major and often only way to foster spatial targeting and obtain a quantitative measure of the required stimulation dose (Bikson et al., 2018). Specifically, the net TMS E-field consists of primary and secondary components. The primary component – the E-field directly induced by the coil – can be determined using scalar and vector potentials, which are calculated using geometric features of the TMS coil. The secondary component – due to induced charges on tissue interfaces – can either be found using the finite element/finite difference method or the boundary element method, respectively. In this study, we develop a method to improve on the speed of calculating the primary component for detailed models of the coil’s 3D geometry or the coil array’s 3D geometry and for potentially very large numbers of observation points pertinent to high-resolution models.

Thielscher and Kammer (Thielscher and Kammer, 2002) developed a model of the coil in the form of an ensemble of *magnetic dipoles.* Briefly, the area of a loop of current is divided into subareas and the magnetic dipoles are placed perpendicular to the loop area in the centers of the subareas. The dipoles are weighted by the loop current and the subarea sizes. This method has further been used in experimental and theoretical studies (see Nummenmaa et al., 2013; Madsen et al., 2015) and in the well-known, open-source transcranial brain stimulation modeling software SimNIBS (Thielscher et al., 2015; Opitz et al., 2015; Nielsen et al., 2018). The method of magnetic dipoles is closely linked to a reciprocity principle (Heller and van Hulsteyn, 1992; Nummenmaa et al., 2013). This principle allows us to reuse standard MEG boundary element method (BEM) computational tools (Meijs et al., 1989; Hämäläinen et al., 1993; Ferguson et al., 1994; Mosher et al., 1999; Gramfort et al., 2014; Tadel et al., 2011; Gramfort et al., 2010; Stenroos et al., 2007; Stenroos and Sarvas, 2012; Stenroos and Nummenmaa, 2016; Nummenmaa et al., 2013; Opitz et al., 2018; Rahmouni et al., 2018) for modeling the secondary TMS field. However, it has been shown that an adjoint double layer formulation of the BEM (Barnard et al., 1967; Makarov et al., 2016; Rahmouni et al., 2018) does not require using the reciprocity principle and magnetic dipoles (Makarov et al., 2018).

A more native model of the coil conductor is that of an ensemble of elementary straight infinitely thin wire segments – *electric dipoles* (or current dipoles). Both primary magnetic and electric fields can be computed from Biot-Savart law and its modification in terms of the magnetic vector potential. Wire-based coil models of specific geometries and of different complexity have been used for numerical optimization and experimental verification in many past and present TMS-related studies (Miranda et al., 2003; Laakso et al., 2014; Koponen et al., 2015; Nieminen et al., 2015; Koponen et al., 2017; Petrov et al., 2017; Gomez et al., 2018).

The ultimate realistic computer-aided design (CAD) model of a coil of any complexity can be simulated with the finite element method (FEM) intended for quasistatic eddy current problems. Deng et al., 2013 used the FEM package MagNet (Infolytica, Inc., Canada) for a comprehensive focality study of 50 different TMS coil designs. ANSYS Maxwell 3D (Cho et al., 2010; Makarov et al., 2016) or SEMCAD X (Rastogi et al., 2017) can be used as well. Commercial FEM packages may be costly and are not without their own limitations. For example, ANSYS Maxwell 3D tends to be slow and may generate large errors for the electric field in air.

Salinas et al., 2007 and Salinas, et al., 2009 have also used the electric-dipole model, but additionally incorporated conductor width, conductor height, and the number of turns to model geometrically realistic coils. They have written two (fast) computer programs in C++ and interfaced them with MATLAB. The first program (3D coil generator) simulates a coil’s geometry; the second program (E-field generator) performs numerical calculations of the induced E-field in the volume of interest. In these studies, the importance of the detailed 3D coil models was emphasized and quantified. Such an approach is a viable alternative to the FEM since it has a comparable accuracy and could take into account the skin effect by locating wire segments close to a conductor’s surface. However, it may result in a large number of elementary current dipoles.

In this study, we attempt to develop this approach further. First, a rather general generator for the coil geometry in MATLAB is constructed, which outputs a detailed electric-dipole model of a coil conductor and of the entire coil. Simultaneously, an equivalent surface CAD model of the coil is created. Second, we interface the electric-dipole model with the fast multipole method (FMM) accelerator for rapid and accurate computations of electric and magnetic fields due to a very large number of current dipoles and at a very large number of observation points. The FMM introduced by Rokhlin and Greengard (Rokhlin, 1985; Greengard and Rokhlin, 1987) speeds up field computations by many orders of magnitude. The FMM is a FORTAN 90/95 program (Gimbutas and Greengard, 2015) compiled within the MATLAB environment and last updated in Nov. 2017.

## 2. Materials and Methods

### 2.1 Distributed current-filament model of a coil conductor

The volumetric coil conductor is modeled as a set of infinity-thin current filaments or segments shown red in Fig. 1. The filaments are always perpendicular to the centerline of the conductor. A number of the filaments passing through conductor’s cross-section can be arbitrarily large. When an alternating electric current flows in a volumetric conductor, two cases are possible: a Litz-wire conductor with uniform current flow through the cross-section or current flow in a thin skin layer close to copper’s surface. In the first case, the current filaments are nearly uniformly distributed over the conductor’s cross-section as shown in Fig. 1a. In the second case, the current filaments are distributed close to its surface as shown in Fig 1b.

**Fig. 1.**
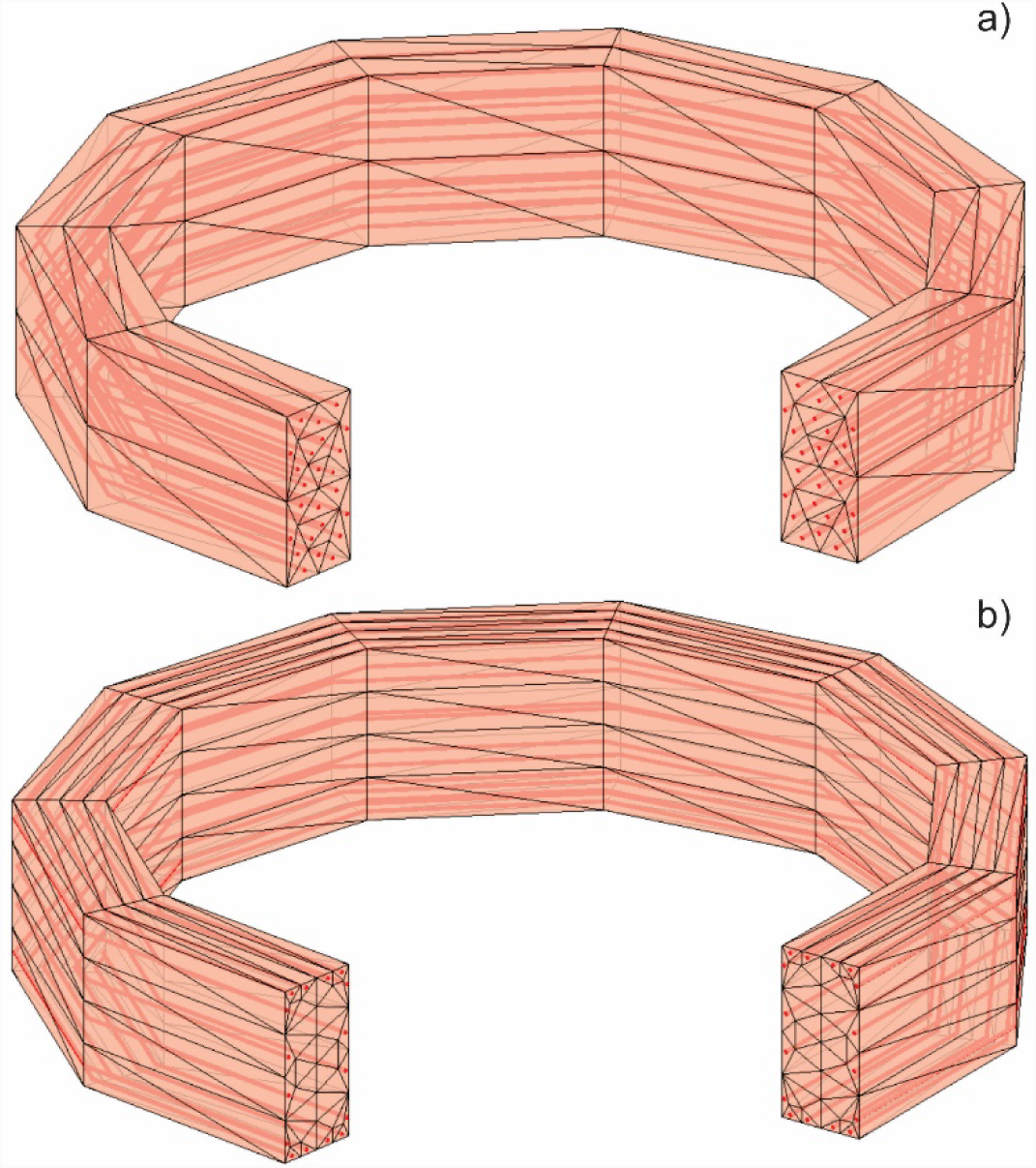
Filaments of current (red) inside conductor’s volume and conductor’s surface CAD model based on triangulation. a) – Uniform current distribution across conductor’s cross-section (Litz wire); b) – modeling the skin effect (a solid conductor at a high frequency).

Consider a small straight element of current *i*_*j*_ (*t*) with segment vector ***s***_*j*_and center ***p***_*j*_. Its magnetic vector potential ***A***^*p*^, at an observation point ***c***_*i*_is given by (Balanis, 2012)

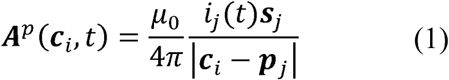

where *π*_0_ is magnetic permeability of vacuum and index *p* means the primary (or “incident” (Balanis, 2012)) field. The electric field generated by this current element is

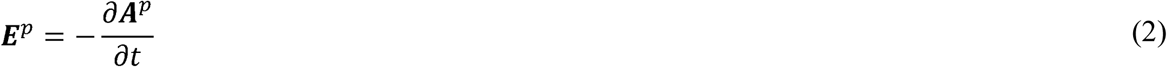

Assuming harmonic excitation with angular frequency *ω* and omitting the phase factor of −*j*, one has

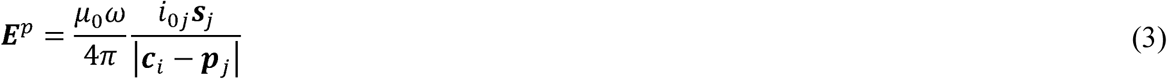

For multiple current segments with moments *i*_*j*0_***s***_*j*_ shown in Fig. 1 and multiple observation points ***c***_*i*_, the net field ***E***^*p*^ at a number of observation points is straightforwardly computed with the FMM as a potential of a single layer repeated three times (Section 2.3).

The same element of current generates a magnetic field ***B***^*p*^ given by the Biot-Savart law

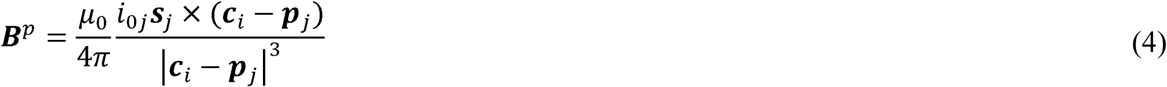

For multiple segments and multiple observation points, Eq. (4) cannot be evaluated using the FMM framework directly. However, it could be rewritten in the form that allows one to use the electric potential of a double layer (layer of electric dipoles) three times (Section 2.3).

### 2.2 Complementary surface CAD model of a coil conductor

A CAD model for the coil conductor is constructed by (i) creating a structured triangular surface mesh for the conductor’s side surface; (ii) creating an unstructured cross-section planar triangular mesh; (iii) creating start/end caps for non-closed conductors; and (iv) joining the meshes and eliminating duplicated nodes. The current filaments in Fig. 1 are defined as short straight lines joining centroids of triangles of the cross-sectional mesh, which is replicated along the conductor’s centerline as many times as necessary. The cross-section is always perpendicular to the conductor’s centerline.

The structured triangular surface mesh for the conductor’s side surface is created based on the array of edges *e* for the conductor’s perimeter (which may either be elliptical or rectangular); the perimeter contour stays the same when moving along the centerline. At every discrete step along the conductor’s centerline (such steps are illustrated in Fig. 1), we add triangular facets in an ordered way. The corresponding MATLAB code accumulates side facets into array *t* as follows

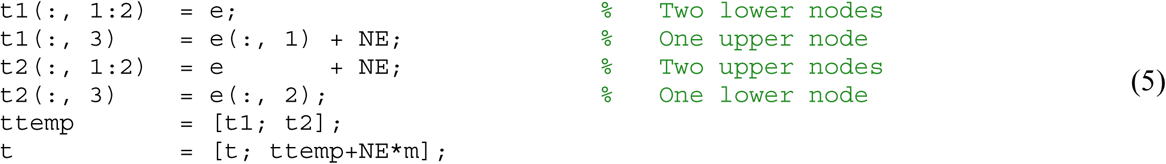

where *NE* is the number of edges (and nodes) for the conductor’s perimeter contour. At every step, we also add *NE* new nodes.

As a result, the solid CAD model of a single coil conductor is created. Then, such a model can be cloned, moved, and deformed as required. Figure 2 shows typical coil models created in this way. Figure 2a is a simplified TMS-MEG coil (Tristan Technologies, San Diego); Fig. 2b is a three-axis multichannel TMS coil array ([22]; Navarro de Lara et al., 2019); Fig. 2c is a simplified MRi-B91 TMSMRI coil model (MagVenture, Denmark) with elliptical conductors of a rectangular crosssection; Fig. 2d is a shapeeight spiral coil with a circular cross-section; Fig. 2e is a double shape-eight spiral coil with an elliptical cross-section and two bootstrapped interconnections. The underlying computational wire grid is always located inside the solid conductor as illustrated in Fig. 1.

**Fig. 2.**
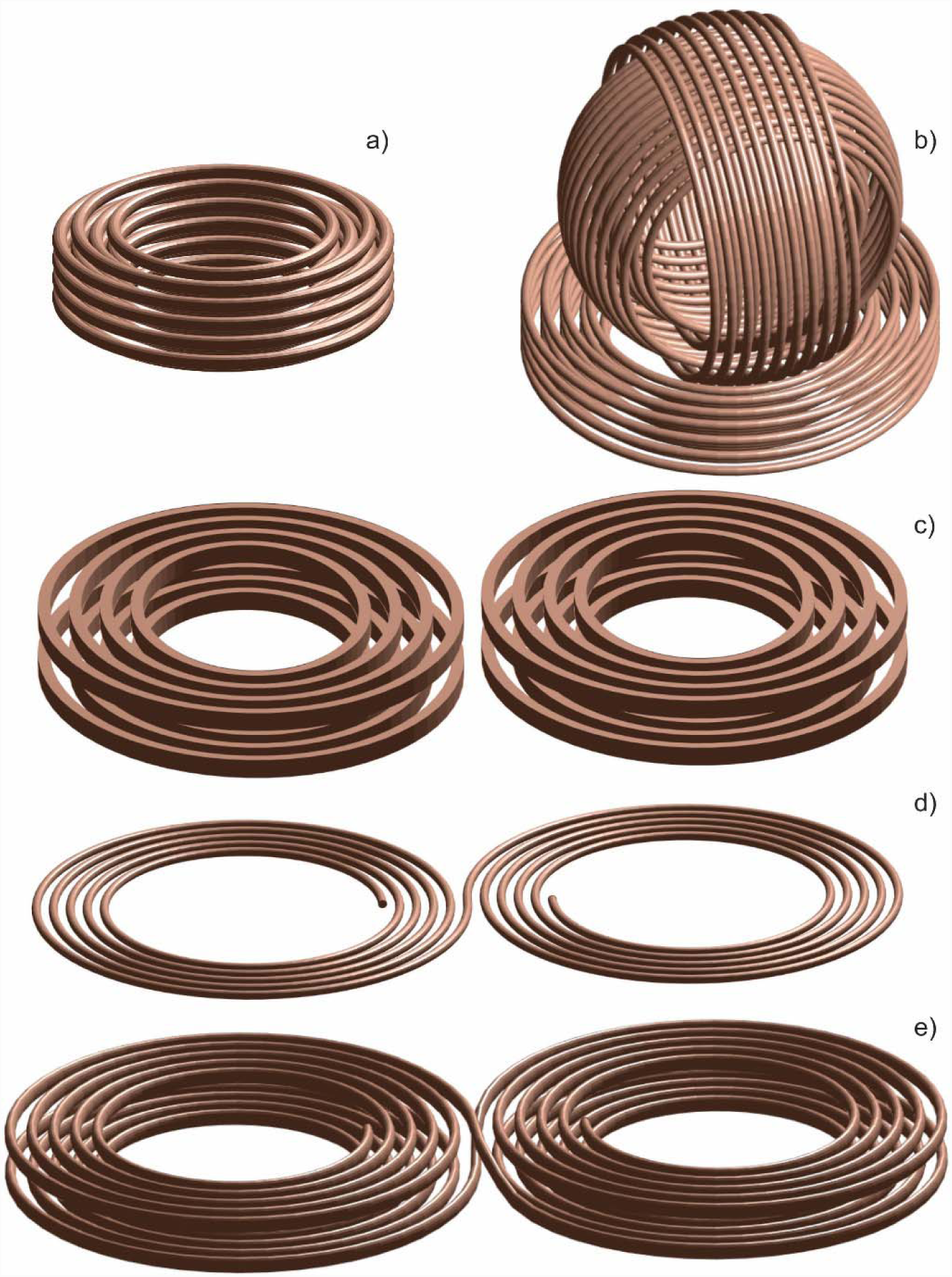
Solid CAD models created using the MATLAB geometry generator. Fig. 2a is a simplified TMS-MEG coil (MagVenture, Denmark); Fig. 2b is a three-axis multichannel TMS coil array ([23]); Fig. 2c is a simplified MRi-B91 TMS-MRI coil model (MagVenture, Denmark) with elliptical conductors of a rectangular cross-section; Fig. 2d is a shape-eight spiral coil with a circular cross-section; Fig. 2e is a double shape-eight spiral coil with an elliptical cross-section and two bootstrapped interconnections.

Once created in MATLAB, the CAD coil model can be exported to any other computational package in STL format.

### 2.3 Fast multipole method for computing *E*^*p*^ and *B*^*p*^

The fast multipole method introduced by Rokhlin and Greengard (Rokhlin, 1985; Greengard and Rokhlin, 1987) speeds up computation of a matrix-vector product by many orders of magnitude. Such a matrix-vector product naturally appears when an electric and/or magnetic field from many current dipoles with moments *i*_*j*0_***s***_*j*_ in Eqs. (3) and (4) has to be computed at many observation or target points ***c***_*i*_.

We use the proven FMM algorithm (Gimbutas and Greengard, 2015) compiled within the MATLAB environment and last updated by the authors in Nov. 2017. In this algorithm, there is no a priori limit on the number of levels of the FMM tree, although after about thirty levels, there may be floating point issues (L. Greengard, private communication). The required number of levels is determined by a maximum permissible least-squares computational error or by the method tolerance, which is specified by the user. The FMM is a FORTAN 90/95 program compiled for MATLAB. In the fastest scenario, the tolerance level `iprec` of the FMM algorithm is set at 0 (the relative least-squares error is guaranteed not to exceed 0.5%).

The electric field in Eq. (3) can be computed as a potential of a single layer repeated three times, i.e., for each component of the field separately. The corresponding pseudo charges used in the FMM function `lfmm3dpart` are to be chosen in the form

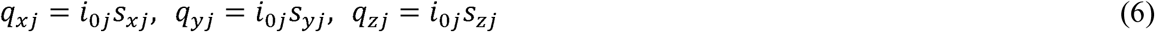

The magnetic field in Eq. (4) cannot be evaluated using the FMM directly. However, it could be

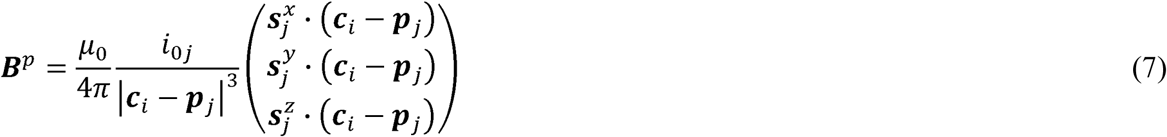

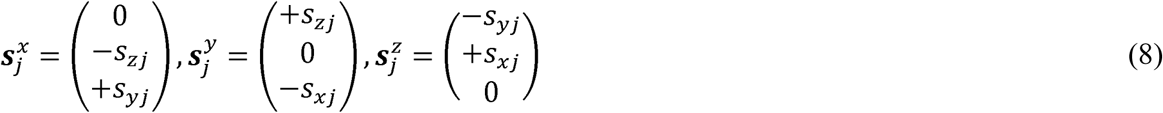

which is equivalent to the electric potential of a double layer (layer of electric dipoles). This potential is to be computed three times and with three different sets of pseudo dipole moments given by Eq. (8) and used in the FMM function `lfmm3dpart` This is another standard FMM task.

### 3. Results

### 3.1 Method accuracy

For an infinitely thin loop of current with the radius of 1 m, the only component of the magnetic field at its axis (–1 m < *z* < 1 m) computed via FMM has been compared with the analytical solution. Given the tolerance level `iprec` of the FMM algorithm equal to 0 and 200 elementary current segments, the relative least squares error is equal to 2.3 x 10^-4^.

As a next example for the same FMM tolerance level, the magnitude of the magnetic field for the MRi-B91 TMS-MRI MagVenture coil model shown in Fig. 2c has been computed via FMM on an observation line whose projection connects the centers of two half windings. The line is located parallel to the coil and rather close to coil’s bottom, at the distance of 5 mm from the bottom conductor. Line length 200 mm is while the coil length is 155mm.

Simultaneously, the same CAD model has been exported into ANSYS Maxwell 3D FEM software and simulated using an FEM eddy current solver (ANSYS 2018 Electromagnetic Suite 19.1.0) with adaptive mesh refinement. The CW excitation frequency is 5 kHz, the coil material is copper, and the skin depth of 0.9 mm is comparable to conductor’s size (3.5mm x 2.2 mm). When the FEM mesh size reaches 0.32 M tetrahedra, the relative least squares error between the two solutions becomes 1.0 x 10^-2^. When the FEM mesh size reaches 1.25 M tetrahedra, the error value does not change significantly. In the FMM model, we have used the skin-depth based current distribution (shown in Fig. 1b) across the conductor’s cross-section, which better accounts for the realistic current distribution.

We were unable to obtain reliable results for ***E***^*p*^ in air using ANSYS Maxwell 3D. Therefore, the FMM solution has been compared against the direct computation given by Eq. (3). For the above example, the relative least squares error for the magnitude of the electric field is 7.4 x 10“^6^.

### 3.2 Method speed

Table 1 reports typical timing results for ***E***^*p*^ and ***B***^*p*^ computations. As an example, a conical-shape coil with 50 single coaxial loops of a circular cross-section has been considered. Each of the loops is divided into 100 straight filaments. We assume a Litz-wire conductor and introduce 30 interpolation points nearly uniformly distributed over the conductor’s cross-section. This results in a coil model that contains 150,000 elementary current dipoles in total.

**Table 1.**
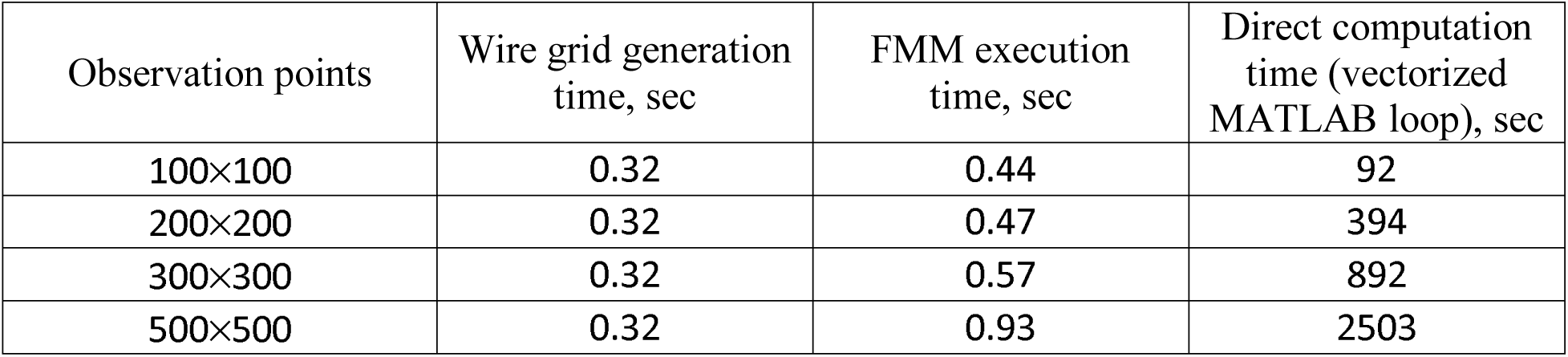
Timing results for ***E***^*p*^ computations. A coil with 50 single loops, each of which is divided into 100 straight cylinders, and with 30 interpolation points per conductor cross-section is considered.

We now keep this coil geometry fixed but vary the number of observation points in a coronal plane whose size is twice the coil size as per Tables 1 and 2. In these tables, we present mesh generation time for the wire grid, FMM execution times, and execution times for a plain yet vectorized MATLAB code which directly computes Eq. (3) and Eq. (4) for the observation points. All results have been averaged over several runs. Here and below in the text, computations are performed using an Intel Xeon E5-2698 v4 CPU 2.2 GHz server, Windows Server 2016 operating system, and MATLAB 2018a.

**Table 2.** Timing results for ***B***^*p*^ computations. A coil with 50 single loops, each of which is divided into 100 straight cylinders, and with 30 interpolation points per conductor cross-section is considered.

| Observation points | Wire grid generation time, sec | FMM execution time, sec | Direct computation time (vectorized MATLAB loop), sec |
| --- | --- | --- | --- |
| 100×100 | 0.31 | 0.49 | 242 |
| 200×200 | 0.31 | 0.52 | 968 |
| 300×300 | 0.31 | 0.62 | 2250 |
| 500×500 | 0.31 | 0.99 | 6320 |

Analyzing the representative FMM computations and the underlying wire model generation times we observe that coil evaluation can generally be performed in less than 1 sec on an ordinary server. On the other hand, the direct vectorized MATLAB loop would run 200-6,000 times slower. We also observe that the FMM code indicates only a very modest increase in time for the reported size of the coil model and the sizes of the observation domain. No specific effort to parallelize the algorithm has yet been made. However, MATLAB automatically performs multithreading pertinent to linear algebra operations available in LAPACK (Linear Algebra PACKage) and some level-3 BLAS (Basic Linear Algebra Subprograms) matrix operations, allowing them to execute faster on multicore-enabled machines.

### 3.3 Evaluation of focality as a function of the number of turns

In this example, some focality characteristics (for more detailed definitions of coil focality see Deng et al., 2013; Koponen et al., 2015; Koponen et al., 2017) of a TMS coil in a coronal plane are investigated. The case in point is a 16 cm long spiral shape-eight coil with two layers and with a rectangular conductor cross-section, shown in Fig. 3a,b. When the number of full turns changes from 2 (as in Fig. 3a) to 5 (as in Fig, 3b), the focality characteristics and the absolute penetration depth change. The goal is to plot the depth, *d*, and the area, *A* of a “circular” segment in Fig. 3c where the electric-field intensity (magnitude of the electric field) exceeds 100 V/m, as a function of the number of turns. One constraint is that the segment domain must be located 20 mm below the coil as in Fig. 3c. The coil current is 7 kA and a CW excitation at 3 kHz is assumed. We also assume that the skin effect is in place.

**Fig. 3.**
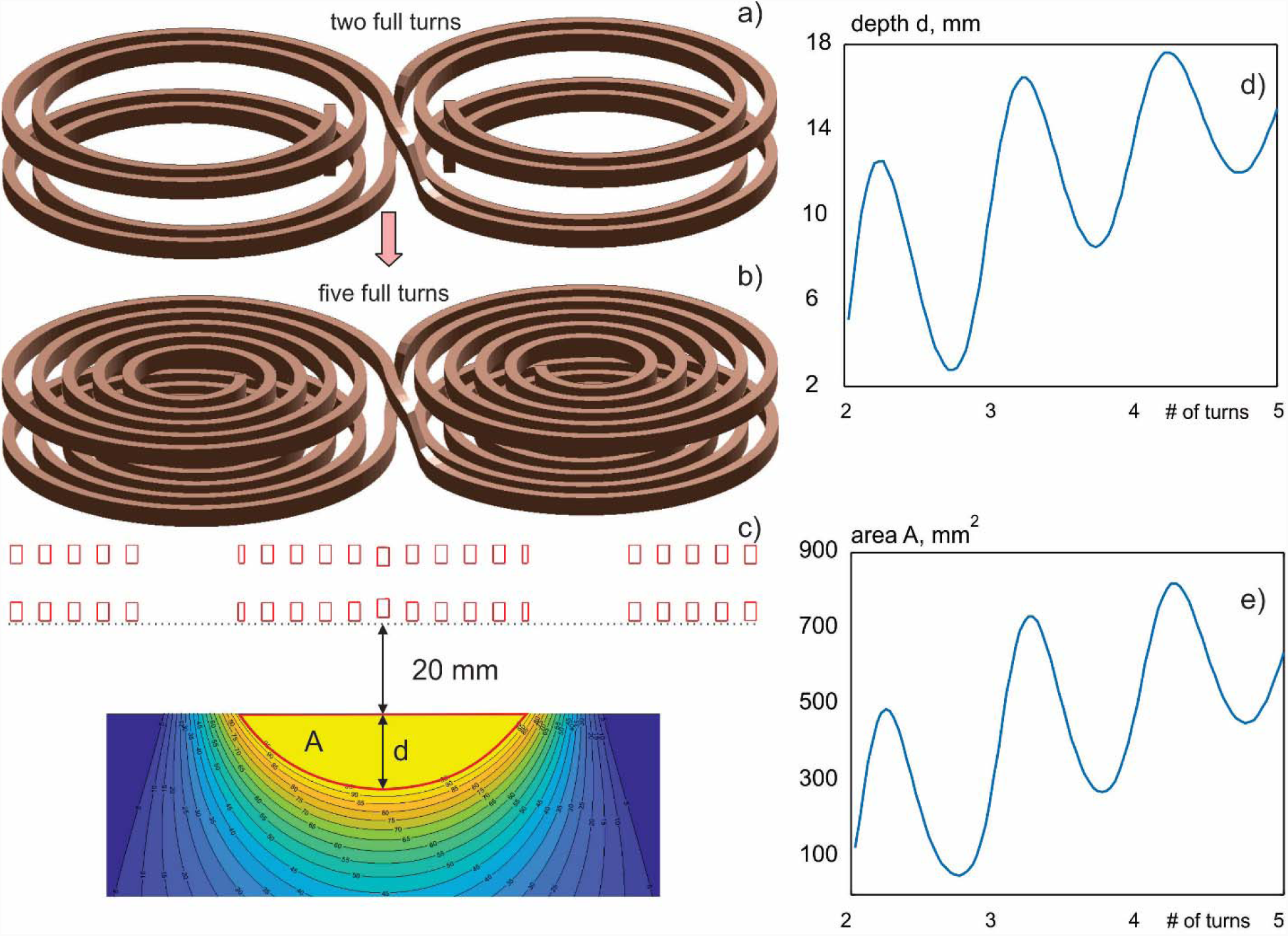
a, b) – Varying coil geometry; c) – focality characteristics in the coronal plane subject to the coil separation constraint; d) – depth of the area under interest as a function of the number of turns; e) – area under interest as a function of the number of turns.

To accomplish this task, an accurate coil model, shown in Fig. 3a,b, has been created for each number of turns with a maximum of 96,000 segments employed for five turns. The number of (non-full) turns is a real non-integer number varying from 2 to 5 in 100 steps. Further, FMM-based field computations at every step are performed for 400×400 observation points covering the domain shown in Fig. 3c. As a result, the curves shown in Fig. 3d (depth of the area under interest as a function of the number of turns) and in Fig. 3e (area under interest as a function of the number of turns) are created. The total run time of the corresponding MATLAB loop is 55 seconds; this time includes generation of 100 wire-based coil models and FMM field computations for every model.

Apart from the timing result, one notes an oscillatory behavior of the penetration depth in Fig. 3d and of the corresponding area in Fig. 3e. This is due to the effect of the farthest side of the coil. This effect may be significant for coils with a relatively small number of turns.

### 3.4 Evaluation of steering in a multi-channel array

The three-axis TMS coil shown in Fig. 2b has been investigated and optimized numerically using the present method. The results have been compared with experimental measurements and reported elsewhere (Navarro de Lara et al., 2019).

In this example, the potential of the three-axis TMS coil shown in Fig. 2b for steering a given polarization is quantified. This coil with the size of approximately 5 × 5 × 5 cm is in fact an array of three coils fed independently, through three synchronized channels. When the x-coil (whose axis is parallel to the x-axis) is excited, its dominant electric-field component 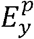 may be altered by the z-coil. Depending on relative channel weighting, the beam steering shown in Fig. 4 appears. The goal is to find the weighting coefficients to enable precise beam steering in the [–2, 2] cm range. The coil current is 5 kA and a CW excitation at 3 kHz is assumed. The coil is made of Litz wire. One constraint is that the field maximum of 120 V/m must occur exactly 2 cm below the coil as in Fig. 4 and at the precise locations along the x-axis.

**Fig. 4.**
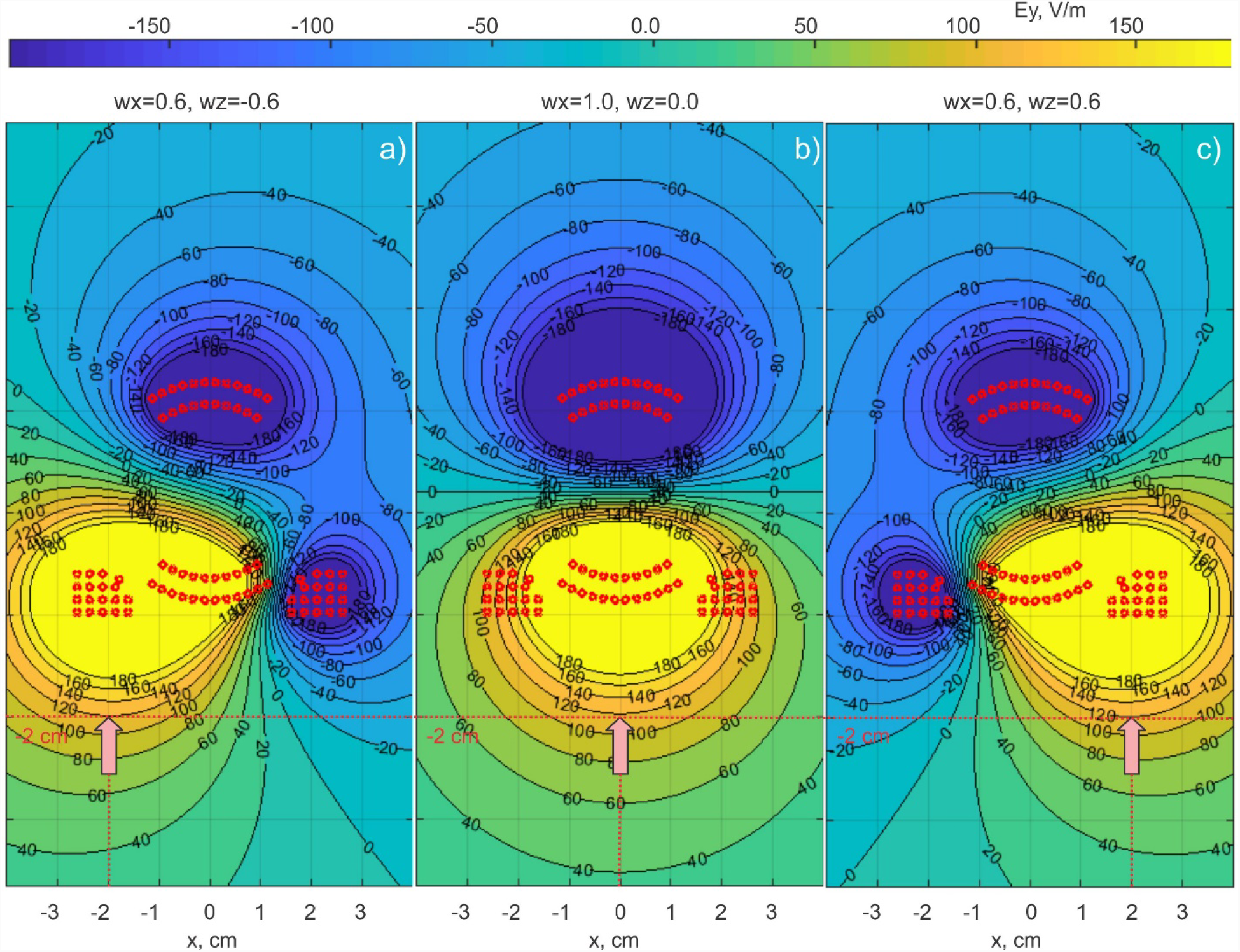
a,b,c) – Beam steering of the dominant electric-field field component as a function of relative weighting of x- and z-coils for the three-axis TIMS coil.

In Fig. 2b) has been created. Further, FMM-based field computations at every step are performed for 400×400 observation points and for different weighting coefficients covering the domain shown in Fig. 4. As a result, the weighting coefficients *w*_*x*_ *=* [0.6, 0.9, 1.0, 0.9, 0.6]; *w*_*z*_ *=* [–0.6, –0.3, 0.0, 0.3, 0.6] have been established to enable constrained steering for *x* = −2,−1,0,1,2 cm as well as for nine intermediate values with the step of 2.5 mm (using symmetry). The CPU run time of one trial is 1.4 sec; this time includes generation of the wire-based coil model and the subsequent FMM field computations. Additionally, approximately 2 seconds are necessary for MATLAB graphics.

Apart from the timing result, one notes that the same steering can be performed in the yz-plane by using the y- and z-coils. Furthermore, steering with x- and y-coils alone is possible.

## 4. Discussion and Conclusion

In this study, an accurate TMS-coil modeling approach based on the coil conductor’s cross-section representation with many distributed current filaments (Salinas et al., 2007) and the use of an efficient FMM accelerator (Gimbutas and Greengard, 2015) was developed and tested. Uniform (Litz wire) or skin-effect based current distributions are included into consideration. The demonstrated speed and calculated accuracy as well as the two application examples provided indicate that this approach is potentially capable of rapid and accurate evaluation of various detailed TMS coil models and arrays of such coils, including their analysis and design.

The MATLAB-based wire and CAD mesh generator for the coil geometry is interfaced with the FMM FORTAN program, which is also compiled within the MATLAB shell. No extra MATLAB toolboxes are necessary. The CAD model of the coil can be imported into other computational software packages in STL format.

The algorithm described above is organized in the form of a MATLAB-based toolkit (TMS Core Lab, MGH). First, a coil model (both wire and CAD) is generated using a dedicated MATLAB script, with one script per coil. The bulk of the manual work here is the definition of the conductor’s centerline using either an analytical formula or a set of points in three dimensions. The output is a coil geometry similar to that shown in Fig. 2.

Then, we compute high-resolution 2D contour plots for any component of the electric and/or magnetic field in the coronal, sagittal, and transverse planes via the FMM, similar to the plots shown in Fig. 4. These two scripts – the mesh generator script and the field computation script – may be further combined into one script and augmented with a parametric loop to enable coil analysis and design, similar to Fig. 3. All other computational scripts including the FMM engine do not have to be modified. They are placed in a separate folder and are only accessed in the form of several well-defined functions.

At present, the geometry modeler is restricted to predominantly flat conductor loops or spirals. The H-coils (see, for example, Deng et al., 2013) with sharp conductor bends in all three planes are subject to further investigation, along with the coil inductance calculations.

## Acknowledgements

The authors wish to thank Dr. Leslie Greengard of the Courant Institute of Mathematical Sciences, New York, NY for useful remarks. This work has been partially supported by the National Institutes of Health Grant R01MH111829.

